# Effect of Human microRNA on Dengue Virus Genome: a cross-specie approach

**DOI:** 10.1101/2022.11.11.516157

**Authors:** Anwesha Deb

## Abstract

Dengue virus infection being a menace worldwide particularly in West Bengal demanded investigation of a new therapeutic approach. In our work we have performed in silico analysis of target specificity of miRNA to dengue virus genome. Several human miRNAs like hsa-miR-4758-3p_MIMAT0019904, hsa-miR-6760-3p_MIMAT0027421, hsa-miR-6839-3p_MIMAT0027581, hsa-miR-6842-3p_MIMAT0027587 are found to be directly interacting with dengue virus proteins like Envelope, NS1, NS3 and 3’UTR region. The target proteins of respective dengue proteins were analysed using stringDB and Cytoscape. Analysis of the target protein interaction network revealed ACTB, AKT1, TLR4 as few of the hub genes that can be indirectly regulated by miRNA and dengue genome interaction. Thus our work focuses on finding novel miRNA that are not known to regulate dengue infection till date and proposes a novel therapeutic strategy.

## Introduction

Dengue virus is an infectious agent of flaviviridae family. The vector responsible for such infection is mainly insects such as *Aedes aegypti* and *Aedes albopictus*. The primary symptoms of dengue infection are dengue fever, dengue hemorrhagic fever and dengue shock syndrome ^1^. Based on WHO reports Dengue is a global threat and has been through quite an upsurge from the year 1960 till today. The reason might be increase in population growth rate, low neighborhood planning, inefficient mosquito planning mostly in the tropical and subtropical regions ^2 3 4^. Although dengue is widely prevalent throughout the world the first report of dengue fever as an epidemic was from Kolkata and East coast of India in the year 1963-1964 ^5^.

Dengue virus is a positive single stranded RNA(ssRNA) virus having serotypes like DENV1, DENV2, DENV3, DENV4^6^. The 11 kb genome consists of three structural proteins, seven non structural proteins and 3’ and 5’ untranslated region. The three structural proteins consist of C protein, prM/M protein and E protein while the seven non-structural ones are NS1, NS2A, NS2B, NS3, NS4A, NS4B and NS5 ^7^. E protein is a large cysteine rich structural protein having 495 amino acids mainly responsible for host cell binding and entry ^8 9^. The structural proteins participates in several biochemical activities, viral RNA replication, cell membrane remodelling and antiviral resistance of host defence mechanism^10^. NS1 is initially translocated into the ER membrane through hydrophobic associations followed by incorporation of N linked carbohydrate moeities and dimerization ^11^. NS1 is secreted from the cell as and when the infection subsides thus resulting in host immune response ^10^. The structural proteins NS2A, NS2B, NS3, NS4A and NS4B are primarily membrane binding proteins. The NS5 protein is the highest conserved among all seven structural proteins throughout the span of 4 dengue virus serotypes ^12 13^

Dengue virus is injected into the human body while the mosquitoes feed onto human blood. The viral pathogen is also spilled over to epidermis and dermis where several dendritic cells like Langerhans and keratocytes are infected ^14 15^. Dengue virus enters the cell primarily by clathrin mediated endocytosis where E protein is bound to multiple membrane receptors like heparin sulphate ^16^, dendritic cell-specific intracellular adhesion molecule-3-grabbing non-integrin (DC-SIGN) ^7 17^. The E protein gets converted from dimeric to trimeric state when shifted from alkaline condition of cell surface to acidic condition of endosome. The change in state of E protein is found to be critical for the binding of host membrane (both mammalian and mosquito) to viral envelope ^18 19^.

### miRNA and regulation of DENV

miRNA biosynthesis involves nuclear protein complex often known as micro processor complex which is present in nucleus of animal cells ^20^. This microprocessor complex has ribonuclease-III protein Drosha, DGCR8 (DiGeorge syndrome chromosomal region 8) and several other minor cofactors like DEAD box RNA helicase p68 (DDX5) and p72 (DDX17). The function of microprocessor complex is to process the pre-miRNAs formed by RNA nuclease II into hair-pin structures of 80-90 nucleotides ^21^. The pre-miRNAs after being exported to cytoplasm by exportin5 protein is cleaved to form a 22 nucleotide sequence miR/miR * complex by DICER, a RNAse III protein ^22 23^. miR of miR/miR * complex gets associated with ago protein and miR * is released in the cytoplasm which eventually gets degraded. Mature miRNA attached to the RISC complex performs its post-transcriptional regulation either by exploiting endonuclease activity of ago2 protein or by blocking the translational process^24^.

miRNA can regulate DENV infection in two ways: controlling the host regulating factors and directly by inhibiting the DENV proteins. Production of Interferon Gamma (IFNɣ) is triggered when viral RNA is detected by Toll Like Receptors(TLR) and Rig-1 Like Receptors (RIR) ^25^. Upregulated IFNɣ can further stimulate inferon stimulating factors using JAK-STAT pathway, thus inhibiting specific steps of Dengue lifecycle ^26^. Although DENV is capable of bypassing production of Interferon Gamma, often mammals are seen to further regulate Interferon Gamma pathway using cellular miRNAs (Pedersen et al., 2007) (Schoggins et al., 2011).

Our primary focus is on regulation of DENV infection by direct interaction of human miRNA with DENV proteins (Figure1). Currently we have multiple evidences that miRNAs are binding to mRNAs as well as directly to the viral genomic RNAs ^28 29^. Although earlier it was shown that host miRNAs are binding to 3’ and 5’ UTR regions of viral genome, later on evidences of binding or miRNA with coding region of viral proteins are also seen ^30 31^. miRNA can target positive stranded viral genomic RNA in the same way as it targets host mRNAs because of high similarity between the two. The viral genomic RNAs targeted by host miRNAs can be either degraded by the host RISC complex or it can get stabilised and lead to viral replication ^32^. Here we are going to mostly discuss a probabilistic approach of miRNAs having direct interaction with Dengue viral genome and further implications.

**Figure 1:**
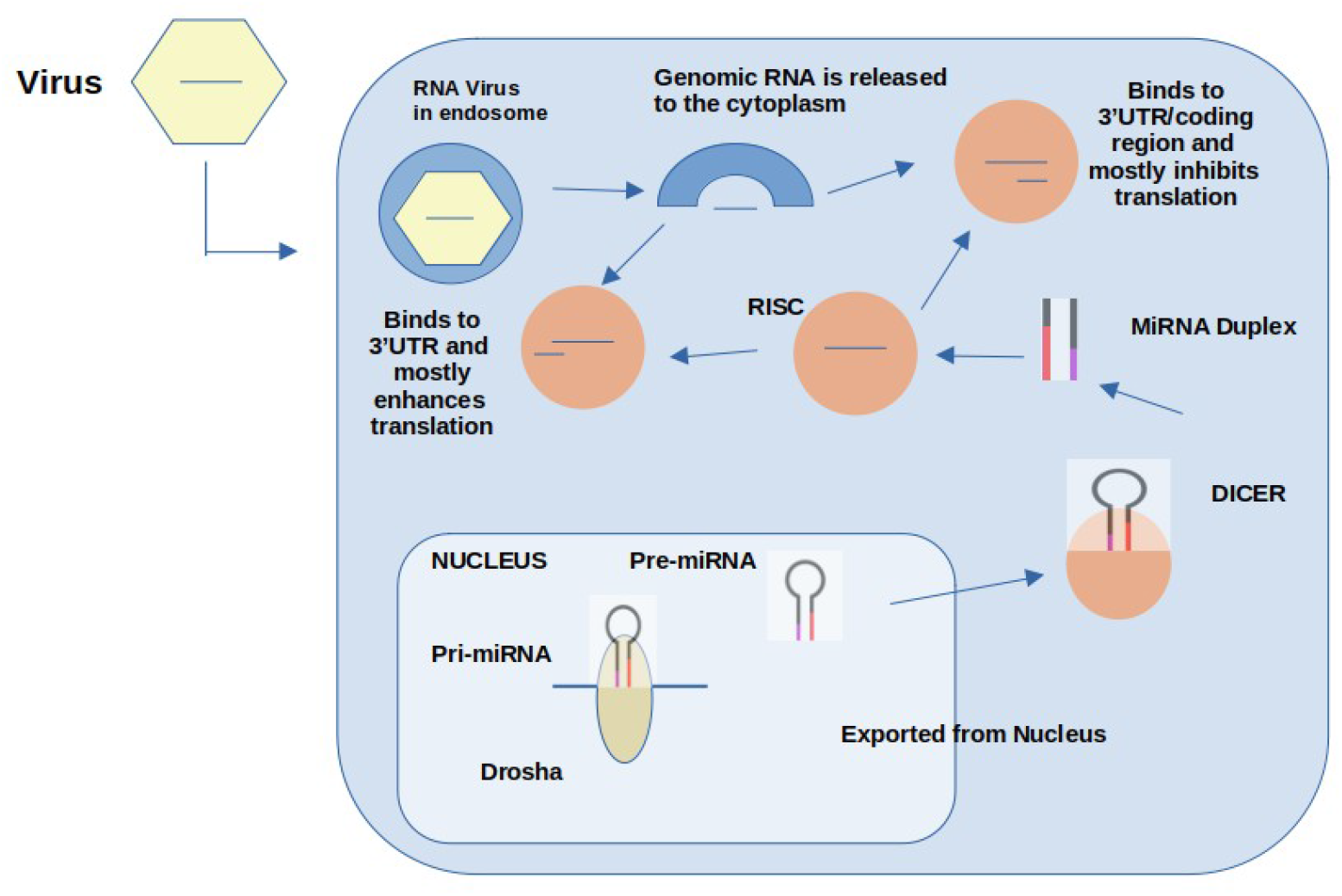
Regulation of DENV infection by direct interaction of human miRNA with DENV proteins

## Materials and Methods

1. **Retrieval of sequences**: Dengue virus genome was retrieved from NCBI genome database and human miRNA sequences were retrieved from mirBase (https://www.mirbase.org/). The data for which is submitted in supplementary ^33^.
2. **Finding target for human miRNA in dengue genome**: Human miRNA having cross species target specificity for dengue genome is assessed using standalone BLAST and psRNATarget webserver (https://www.zhaolab.org/psRNATarget/). The minimum free energy criteria for selection of targeting miRNA was fixed to −18 Kcal ^34^.
3. **Analysis of miRNA target site accessibility**: The target sites selection for miRNA binding is one of the most critical part of predicting miRNA mediated gene suppression. In this paper it has been done manually by taking 150 bps sequence from upstream and downstream of the seed region using UNAFold webserver (http://www.unafold.org/)(Markham & Zuker, 2008).
4. **Gene prediction of miRNA target sites**: miRNA target analysis on the dengue virus genome is detected using RNAhybrid webserver (https://bibiserv.cebitec.uni-bielefeld.de/rnahybrid). The MFE was kept at −18 Kcal/mol for selection of optimal binding.
5. **Genome annotation**: The target site prediction was followed by genome annotation from Virus Pathogen Resource database (https://www.viprbrc.org/brc/home.spg?decorator=vipr). The GATU genome annotator tool is used for indexing the Dengue genome. The gene annotation table was downloaded and manually checked for the target sites of miRNA.
6. **Finding novel miRNA**: The novel human miRNA having cross-specie target specificity are identified from literature study.
7. The analysis of miRNA targeted dengue proteins with human proteins were performed using literature survey and from server DENHUNT(http://proline.biochem.iisc.ernet.in/DenHunt/).

## Result

The mature human miRNA sequences were downloaded from miRBase in fasta format and aligned against dengue genome using blastn suit of standalone blast and also using the automatic webserver psRNATarget. The best candidates among them were selected and checked for target accessibility using UNAfold webserver (http://www.unafold.org/)(Fig:2). In order to ensure specific binding we have kept bulge loop allowance of 0-3 bp and mismatch allowance of 2 bps ^35^. The final 6 miRNAs were taken and aligned against dengue genome using RNA hybrid server. We get the positions at which they are binding and also the MFE value of miRNA binding to the dengue genome. The miRNA corresponding to MFE values below −18 kCal/mole were selected. Further GATU the Genome Annotator tool was used to understand the specific domains to which the miRNAs are binding (Table1)(Fig: 3). After an extensive literature survey on interaction of human miRNAs and dengue genome we came up with four novel miRNAs which in all probabilities are not yet mentioned to be involved in dengue infection. The particular target domains of Dengue are checked for their immunogenic responses in mammalian both from literature and Denhunt database. Further functional analysis of target mammalian proteins are discussed in the next section.

**Figure 2:**
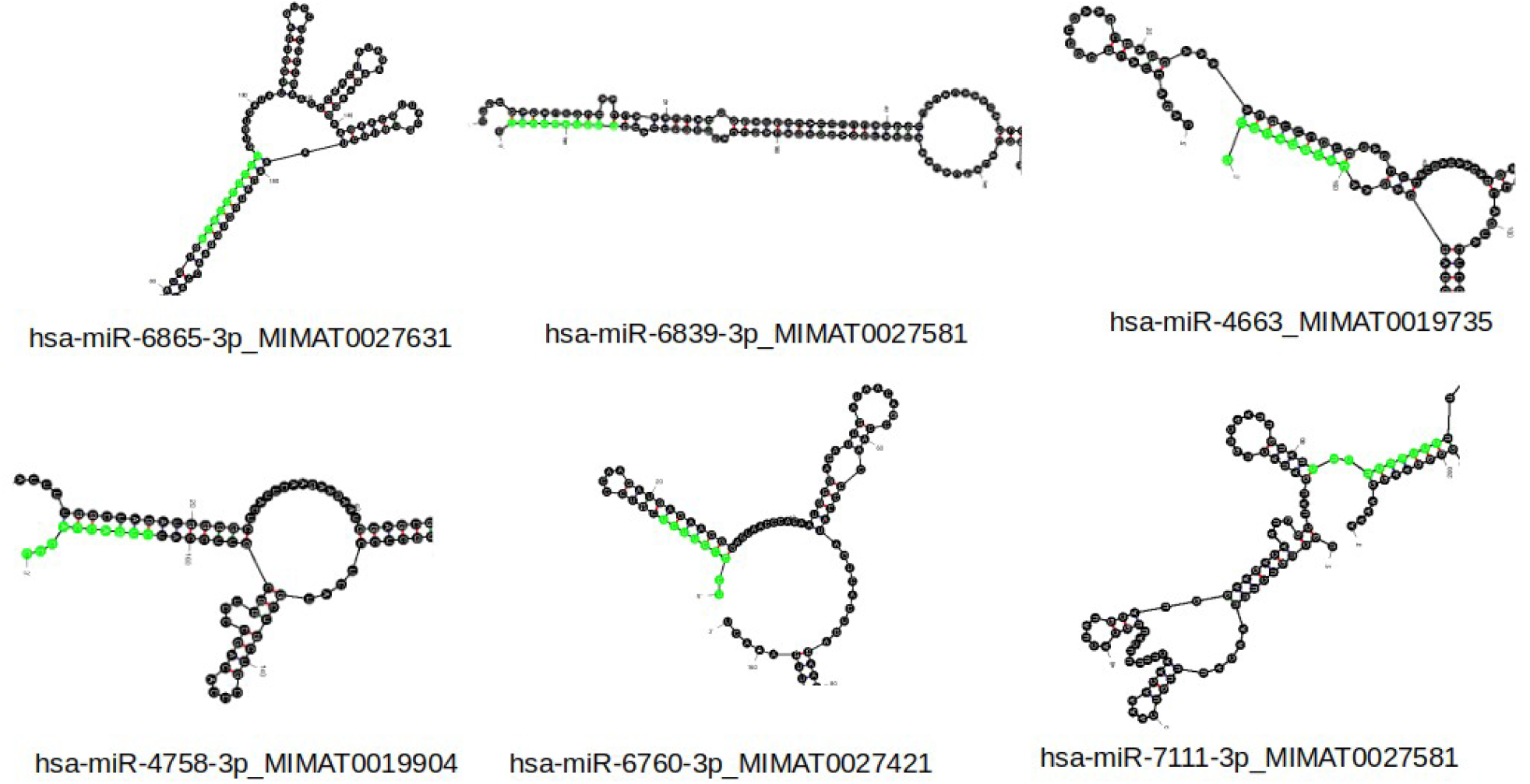
The best miRNAs selected on the basis of target accessibility and the Gibbs Free Energy

**Table 1:** miRNAs predicted to have cross specie specificity and there respective dengue virus target domain. The MFE was set to be −18 Kcal/mol and all interactions having value <= −18 Kcal/mol.

| miRNA | Targeted Dengue protein | Energy in Kcal/mol |
| --- | --- | --- |
| hsa-miR-4663_MIMAT0019735 | NS5, NS1, DENV-Envelope | -23.2 |
| hsa-miR-6865-3p_MIMAT0027631 | NS1 | -23.0 |
| hsa-miR-4758-3p_MIMAT0019904 | NS3 | -25.2 |
| hsa-miR-7111-3p_MIMAT0028120 | NS3 | -25.6 |
| hsa-miR-6839-3p_MIMAT0027581 | 3'UTR | -22.6 |
| hsa-miR-6760-3p_MIMAT0027421 | DENV-Envelope, Protein (Pr) | -24.1 |

**Figure 3:**
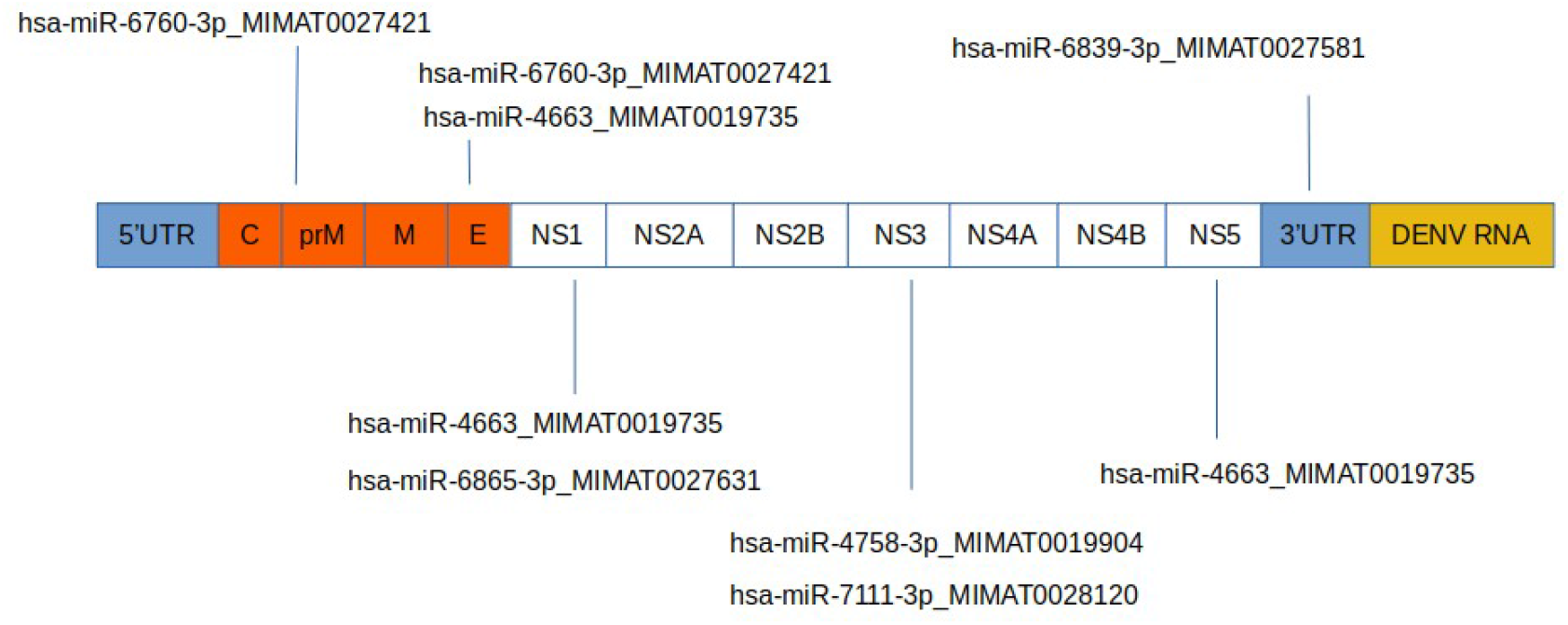
Schematic representation of selected miRNAs and their target domain in dengue genome

## Discussion

hsa-miR-4663_MIMAT0019735 and hsa-miR-6760-3p_MIMAT0027421 are binding to the DENV envelope (Table:1) protein which is known to be primary mediator in attachment of the virus to the host cell ^19,36^. Among two of the identified miRNAs identified having cross specie target specificity for Dengue virus, hsa-miR-6760-3p_MIMAT0027421 is all probabilities is a novel finding in terms of having specificity for dengue virus. Thus hsa-miR-6760-3p_MIMAT0027421 can be further investigated i*n vitro* and *in vivo* systems to reveal its functionality in blocking the envelope protein from binding to the host cell surface receptors. Envelope protein gets attached to the cell surface receptors and is placed on clathrin coated pits further getting transported into endosomes. Once the early endosome is transformed to more acidic late endosome the E protein changes its conformation from homodimer to homotrimer. This homotrimer fuses with membrane of endoplasmic reticulum and releases the viral genomic material followed by its replication ^37^(Fig:4). Since Envelope protein of Dengue virus is of extreme importance we have also performed network analysis using Stringdb and Cytoscape of its target proteins. The network was analysed in Cytoscape and a protein called ACTB was determined to be the hub protein having highest degree and betweenness-centrality values (Fig:5). ACTB is an actin protein which is shown to be critical for endocytosis in Fig:4 and is indeed present in higher quantities in extracellular space, cytoskeleton, cytosol and plasma membrane. All these four compartments are essential for endocytosis of virus particle. In the figure 6 we have an interesting observation that actin protein (ACTB) is expressed in higher amounts in tissues/organs that are more prone to dengue infection ^38^. In organs like liver and skin ACTB expression is very high and are known to be critical for severe dengue infections. The other two organs lungs and kidney are also prone to dengue infection are also having comparatively higher expression of ACTB than many others, although not to the level of Liver and Skin. Thus hsa-miR-4663_MIMAT0019735 and hsa-miR-6760-3p_MIMAT0027421 are potential therapeutic candidates which can be monitored for its ability to downregulate DENV envelope protein.

**Figure 4:**
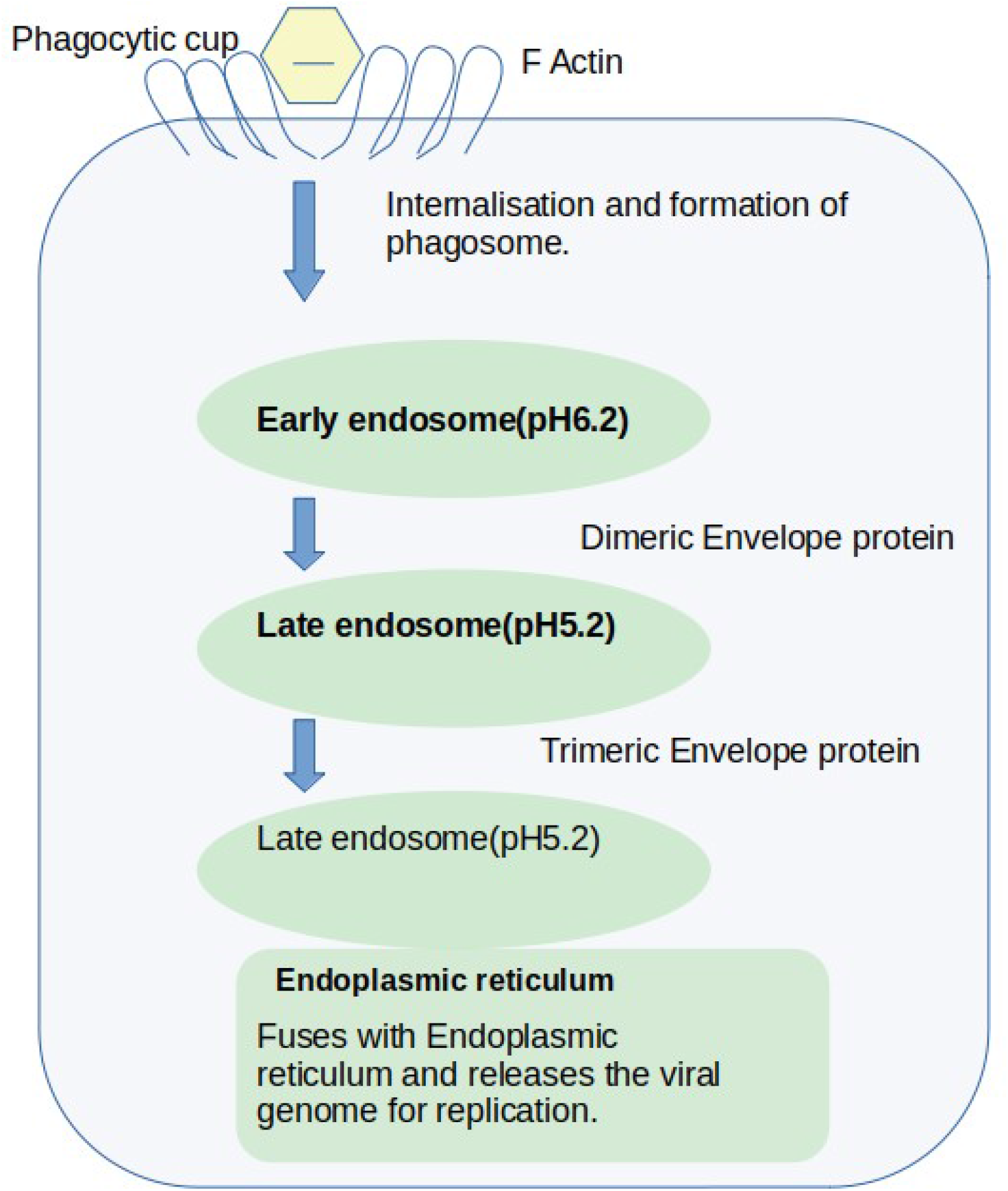
Role of Actin protein in internalisation of viral particle

**Figure 5:**
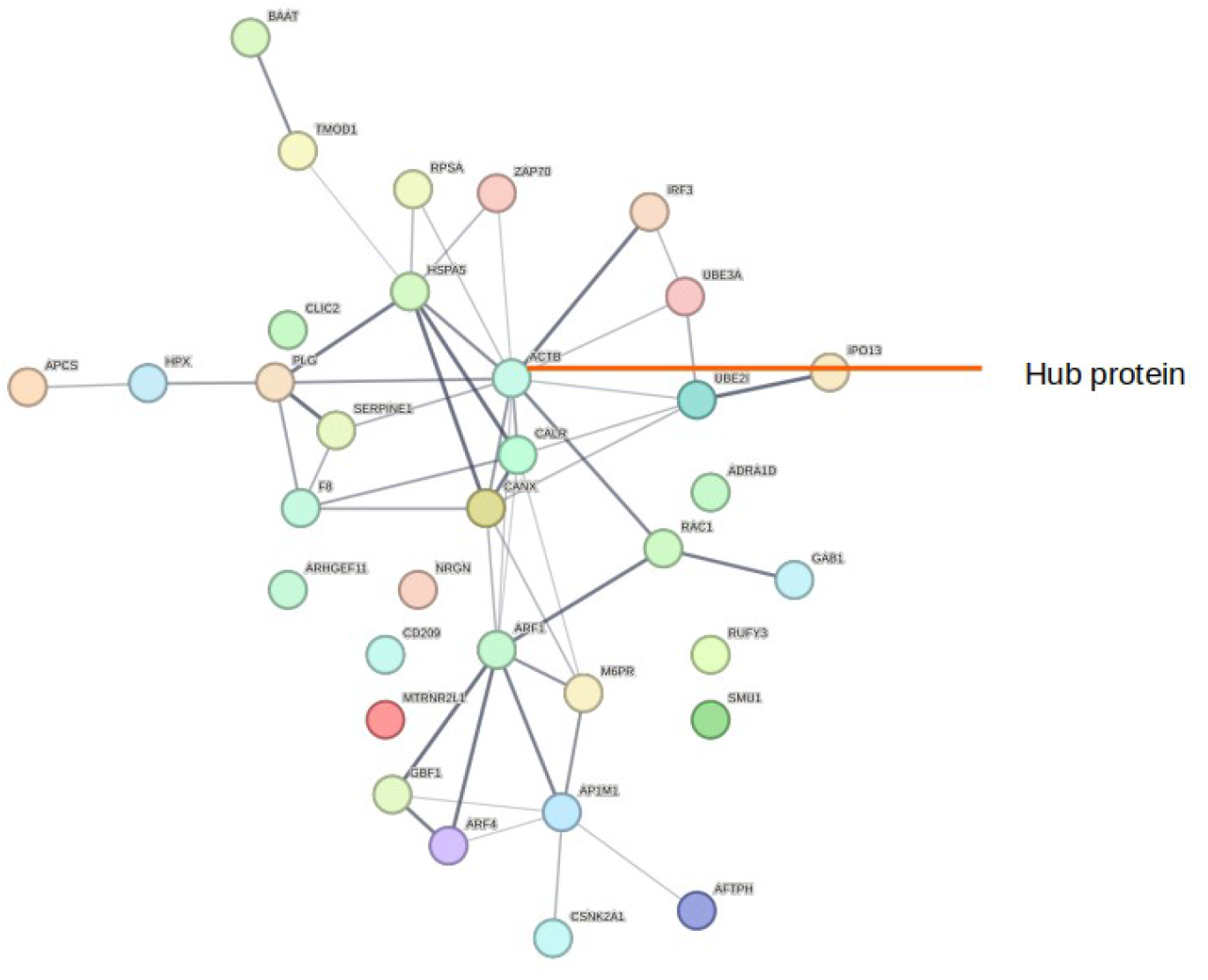
Network analysis of proteins targetted by Dengue Envelope Protein

**Figure 6:**
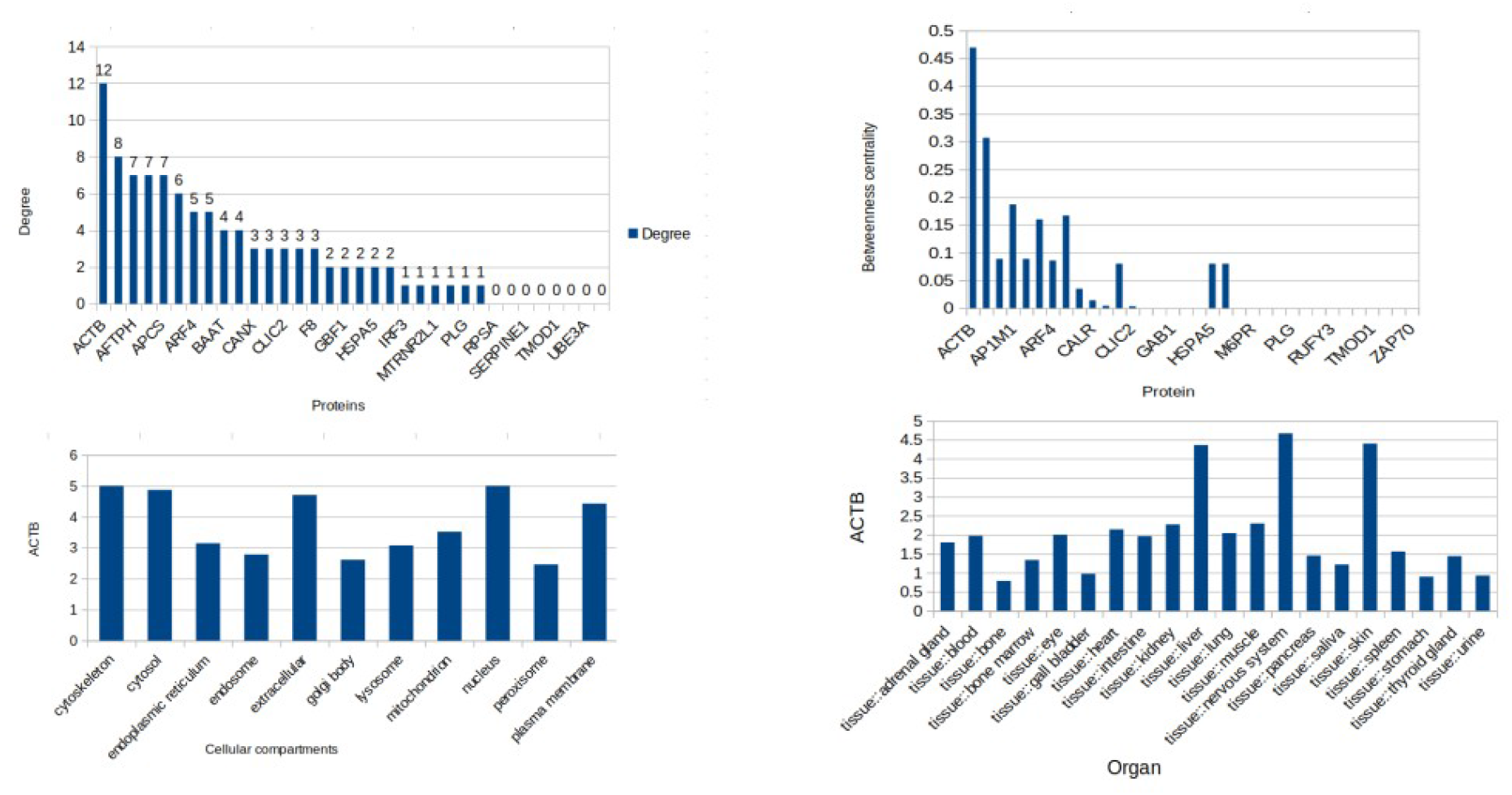
Profiling of hub proteins targetted by Dengue Envelope Protein

hsa-miR-6760-3p_MIMAT0027421 interestingly is also having target specificity for prM protein (Fig:3) of DENV(Table:1). There are several evidences that E protein of dengue virus depends heavily on prM protein for performing few of its functionalities ^39^. prM is also important for stabilisation of temperature sensitive epitope of Envelope protein and when co-expressed with E protein it often stabilises the vaccine. Thus targeting its expression with hsa-miR-6760-3p_MIMAT0027421 can be a therapeutic approach.

hsa-miR-4663_MIMAT0019735, hsa-miR-6865-3p_MIMAT0027631 can be seen targeting NS1 protein of DENV virus which primarily known for activation of complement pathway(Table:1). Network analysis of proteins targeted by NS1 domain showed three primarily important clusters: TLR4 mediated pathway (yellow), AKT1 mediated pathway(green), Ribosome protein cluster(FIG:7). Modhiran et. al. has shown in their paper that NS1 can directly affect TLR4 activated pathway in severe dengue patients(Fig:9) ^40^. Severe dengue cases are often associated with increased levels of TNF-ɑ, IL-1β, IL-6, IL-8, IFN-ɣ and MCP-1 ^41,41^. Increased level of TNF-ɑ, IL-1β, IL-6, IFN-ɣ results in depletion of barrier function of endothelial cells and increased level of MCP1 and IL-8 results in alteration of tight junctions between dengue infected cells. Dengue haemmorhagic fever is mediated by TNF-ɑ by apoptosis of endothelial cells(Fig:9). It has been shown that antibodies antagonistic to TLR-4 were able to reduce NS1 mediated immune response ^40^. Thus we can also take similar approach by inhibiting NS1 domain of Dengue virus and thereby controlling its immune effect.

**Figure 7:**
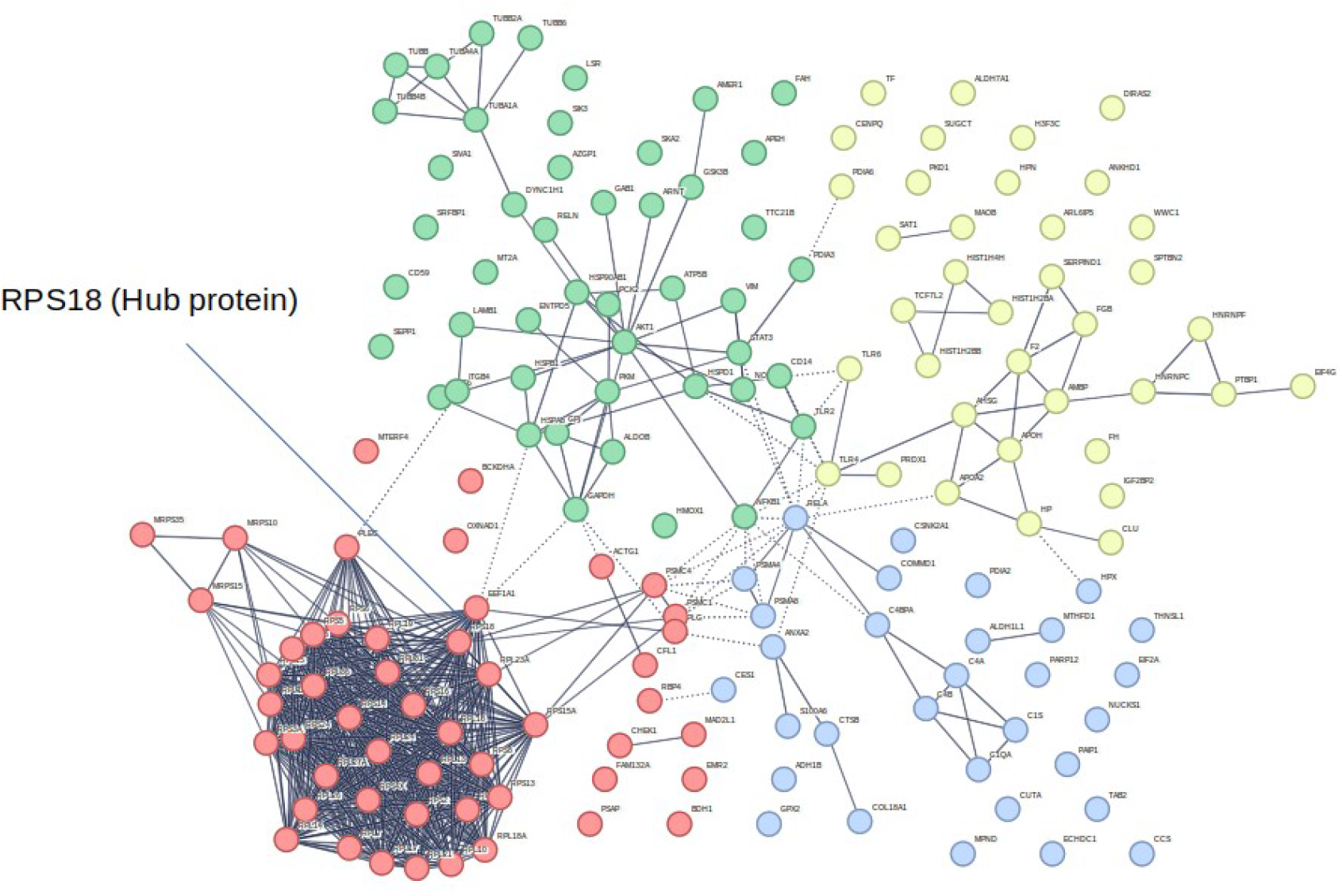
Network analysis of proteins targeted by NS1 domain showed three primarily important clusters: TLR4 mediated pathway (yellow), AKT1 mediated pathway(green), Ribosome protein cluster

**Figure 8:**
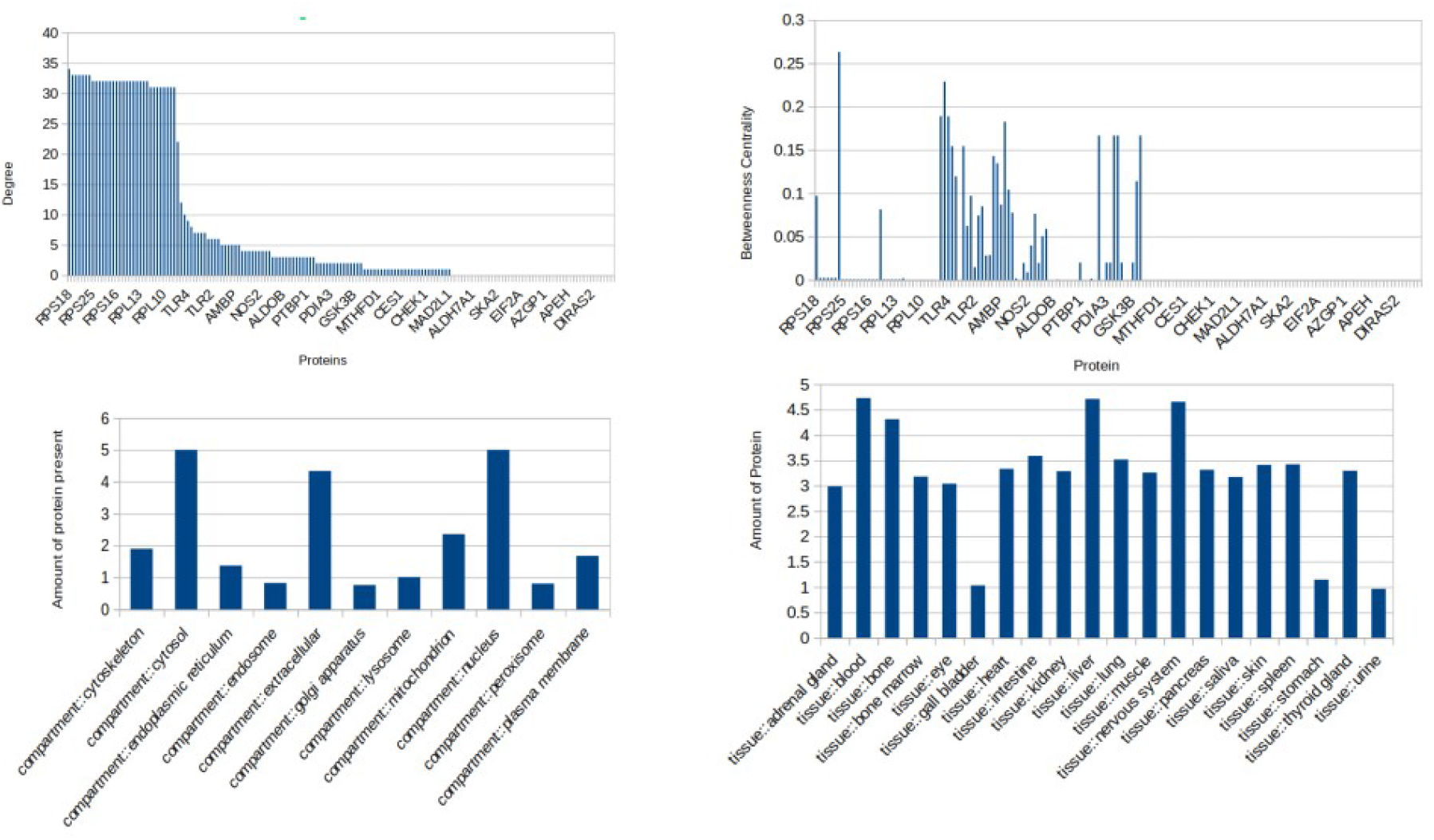
Profiling of hub protein RPS18 targetted by Dengue NS1 Protein

**Figure 9:**
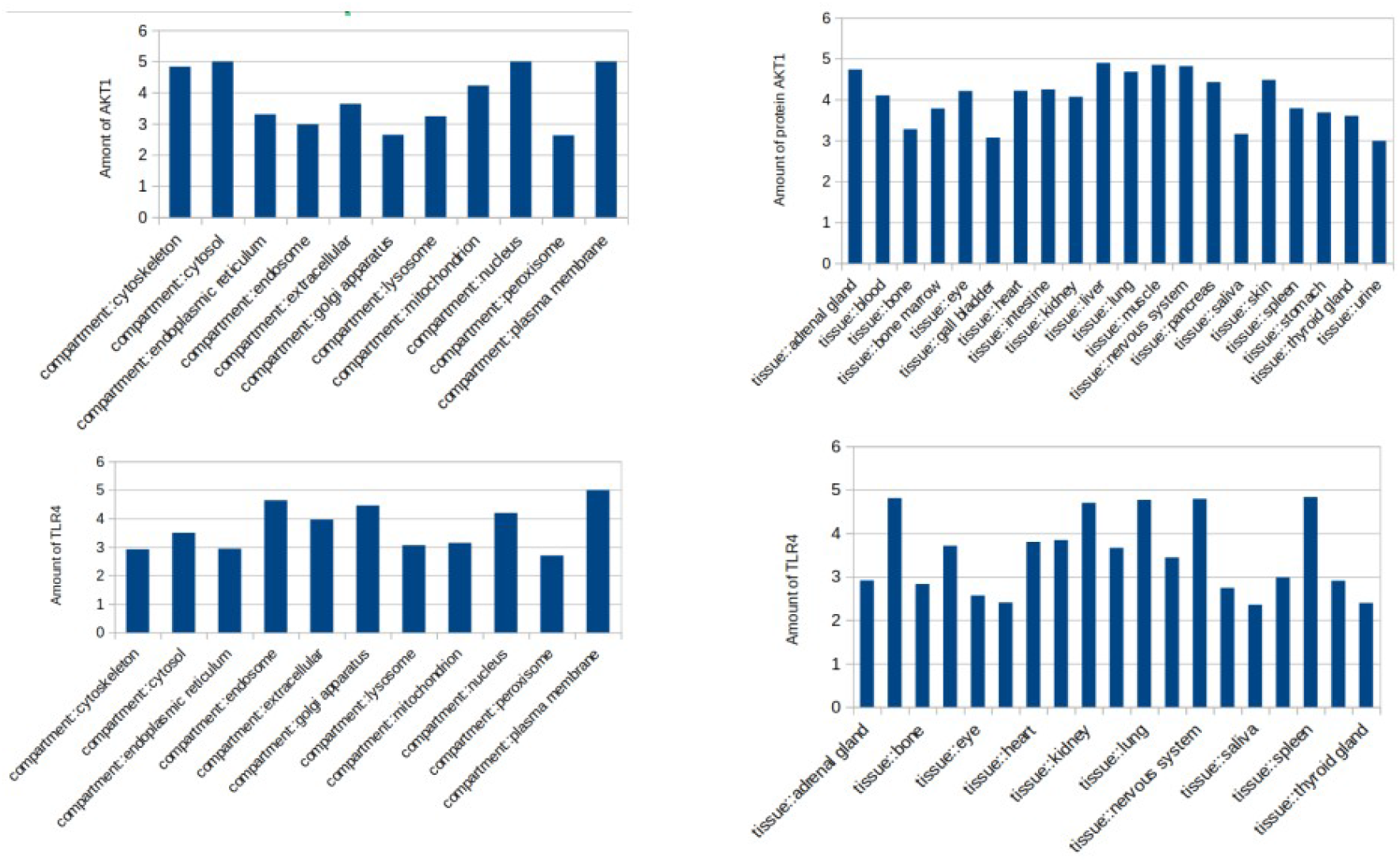
Profiling of hub protein TLR4 targetted by Dengue NS1 Protein

Next cluster consists of AKT1 signal transduction proteins involved in AKT/PI3K/mTOR mediated megakaryopoesis (FIG:7). Cell survival, reduced apoptosis and cell maturation of megakaryocytes is dependent on PI3K/AKT pathway ^42^. Increased presence of phospho-AKT and PI3K has been observed in DENV infection, both of which regulates mTOR. mTOR in turn is responsible for megakaryocyte survival and its transformation to platelets ^43^. Eventually it was noticed that two regulatory subunits of PI3K; p85 and p110ɑ are having reduced expression in DENV infected patients which might result in increased rate of megakaryocyte destruction^44^. The cluster also contains a subcluster of cytoskeleton proteins like actin and tubulin. It has indeed been mentioned that mTOR also regulates ploidy of megakaryocytes by regulating proteins like tubulin thus affecting megakaryopoeisis^44^.

The final and most important cluster of NS1 target protein involves ribosomal protein RPS18(FIG:7). It was observed in multiple cases that 50% of NS1 targeting proteins are ribosomal proteins or per se the whole ribosome ^45^. The ribosomal proteins are also having few extraribosomal functions like ribosomal binding to endoplasmic reticulum, apoptosis, cell cycle etc. along with its translational abilities^46^. Since RPS18 protein expression is not different for normal and infected cells it had to be silenced with siRNA to look for its effects ^45^but it was found that RPS18 forms an important cluster of NS1 targetting proteins. It was found that DENV infection results in relocation of RPS18 protein to the perinuclear position and indeed it is found to have profound effect in virus replication^47^. Involvement of RPS18 protein in DENV replication can also be suggested by its high amount of presence in nucleus compartment (Fig:8).

hsa-miR-4758-3p_MIMAT0019904, hsa-miR-7111-3p_MIMAT0028120 are having specificity to NS3 domain of Dengue virus(Table 1)(Fig:3). The target proteins for NS3 were downloaded from Denhunt and submitted to stringdb for studying the protein interactions. The network was then imported to Cytoscape for further analysis and ACTB was found to be the hub protein (FIG:10, FIG:11). There are several instances in literature where we find that NS3 and NS5 bind to several cytoskeleton proteins like ACTB, ACTG1 etc. Interestingly the literatures are also having mentions of proteins like CEP250 which also has a high betweenness centrality from our analysis(Fig:11) ^48^. High betweenness of CEP250 is indeed justified because it is a protein of kinesin family and can be bound to multiple cytoskeletal framework proteins. Also the ability of flavivirus to disrupt the cytoskeletal framework of a cell is a well known fact, hence justifying our network analysis result ^49^. The other proteins in the network are LMNA which helps in nuclear localisation and NFΚβ signalling proteins which helps in recognition of nuclear localisation signal sequence and thus promoting viral replication. Probably it has been reported for the first time that hsa-miR-4758-3p_MIMAT0019904 is targeting NS3 domain of Dengue Virus. Since NS3 is an important pathogenic component of dengue virus, targeting it with small non-coding RNAs can be a critical therapeutic strategy.

**Figure 10:**
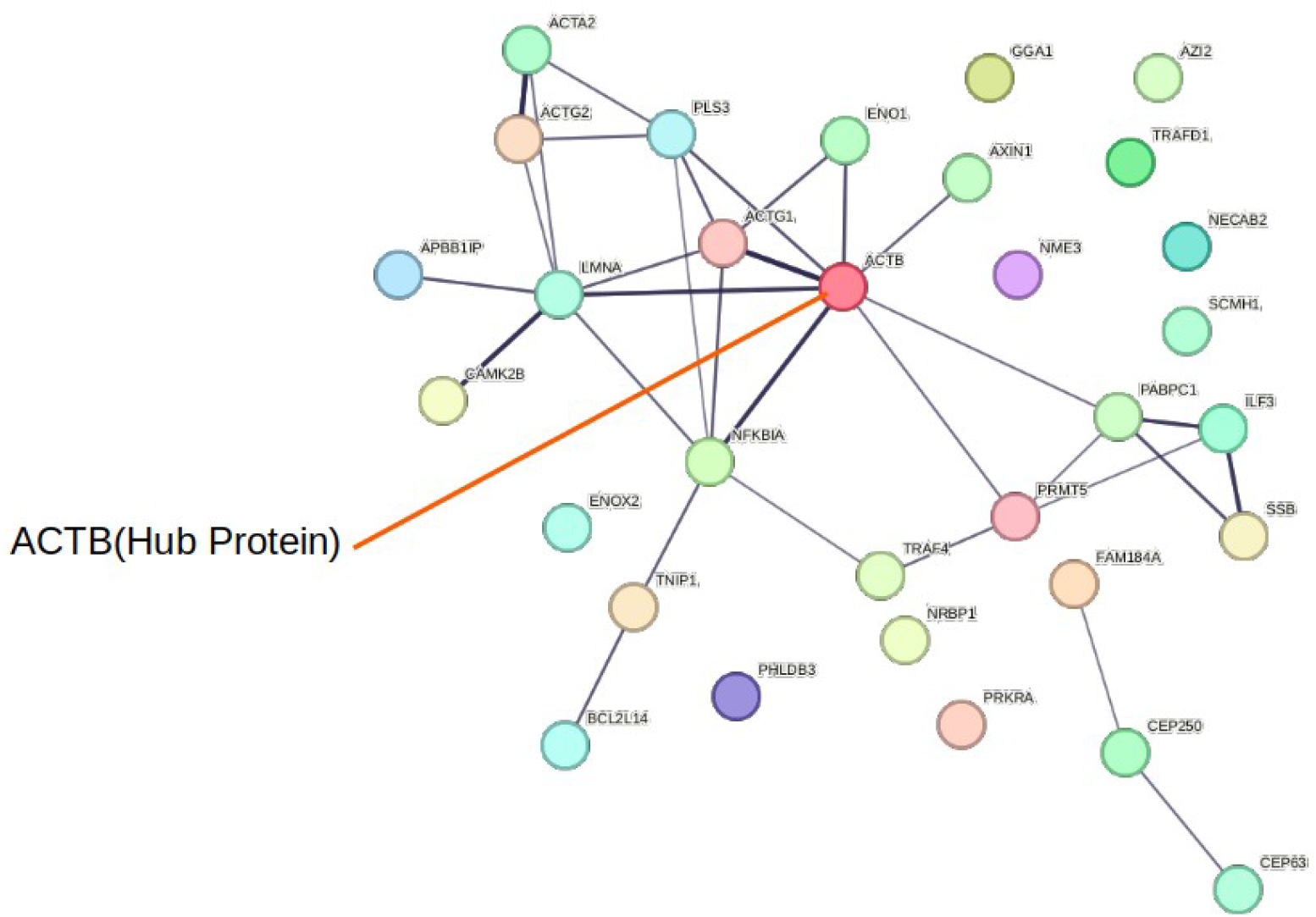
Network analysis of proteins targeted by NS3 domain

**Figure 11:**
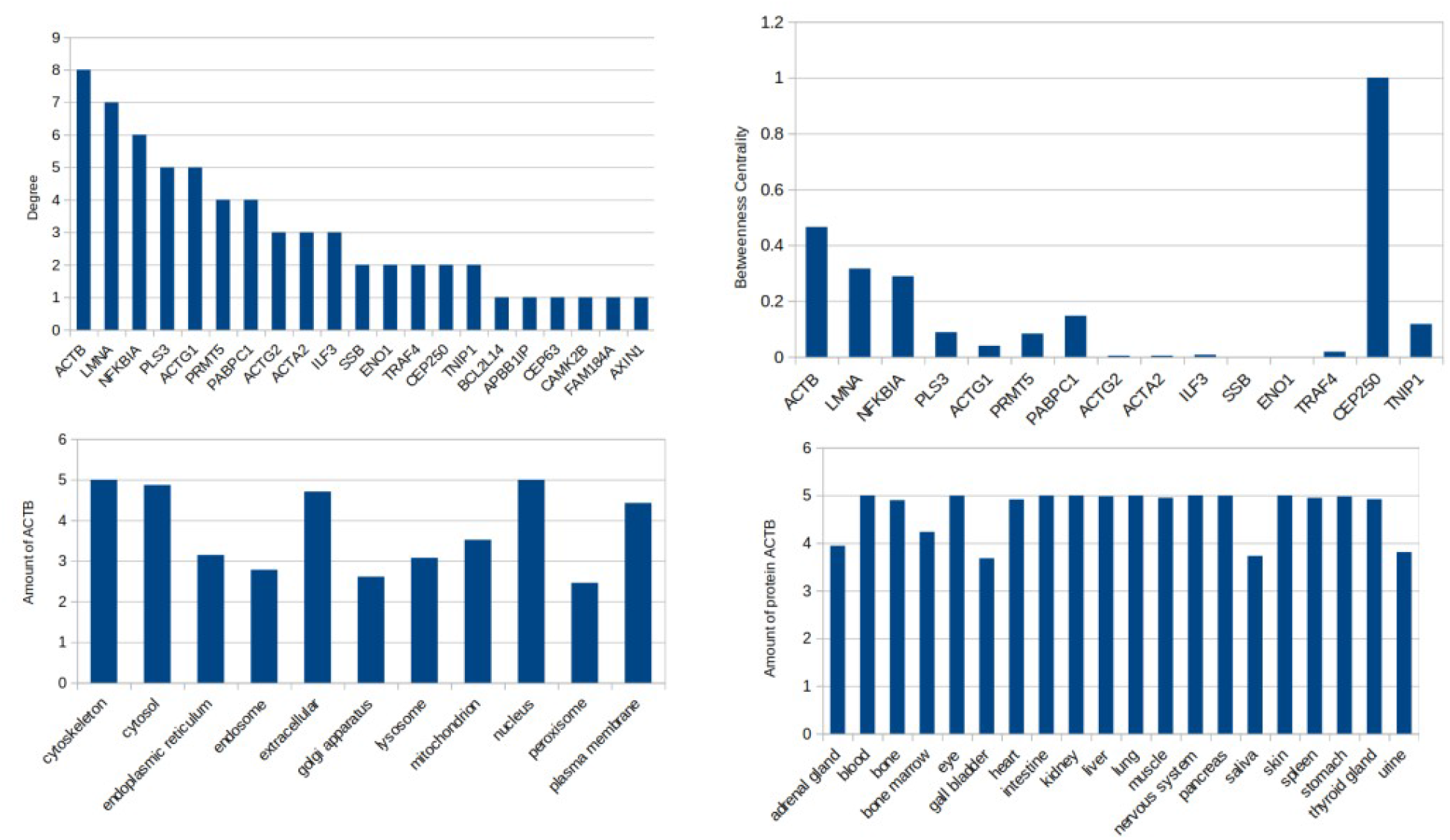
Profiling of hub protein ACTB targetted by Dengue NS3 Protein

hsa-miR-6839-3p_MIMAT0027581 is observed to be bound to 3’UTR domain of Dengue virus which is again a critical domain in mRNA translation of viral genome. Till date no literature has mentioned hsa-miR-6839-3p_MIMAT0027581’s involvement in dengue infection, which makes it a more interesting finding. The target proteins of 3’UTR region consists of several transcriptional and translational regulators among which YBX1 is the hub protein(Fig:12). It is a known fact that dengue RNA translation can be regulated by binding of YBX1 protein throughout the genome ^50^. 3’UTR region of dengue virus has binding region for YBX1 which is involved in translational regulation^51^. Therefore it can be presumed that YBX1 is having a direct role in upregulation of viral genome replication. Blocking of 3’UTR with hsa-miR-6839-3p_MIMAT0027581 might result in reduced binding of YBX1 to 3’UTR, thus reduced viral replication.

**Figure 12:**
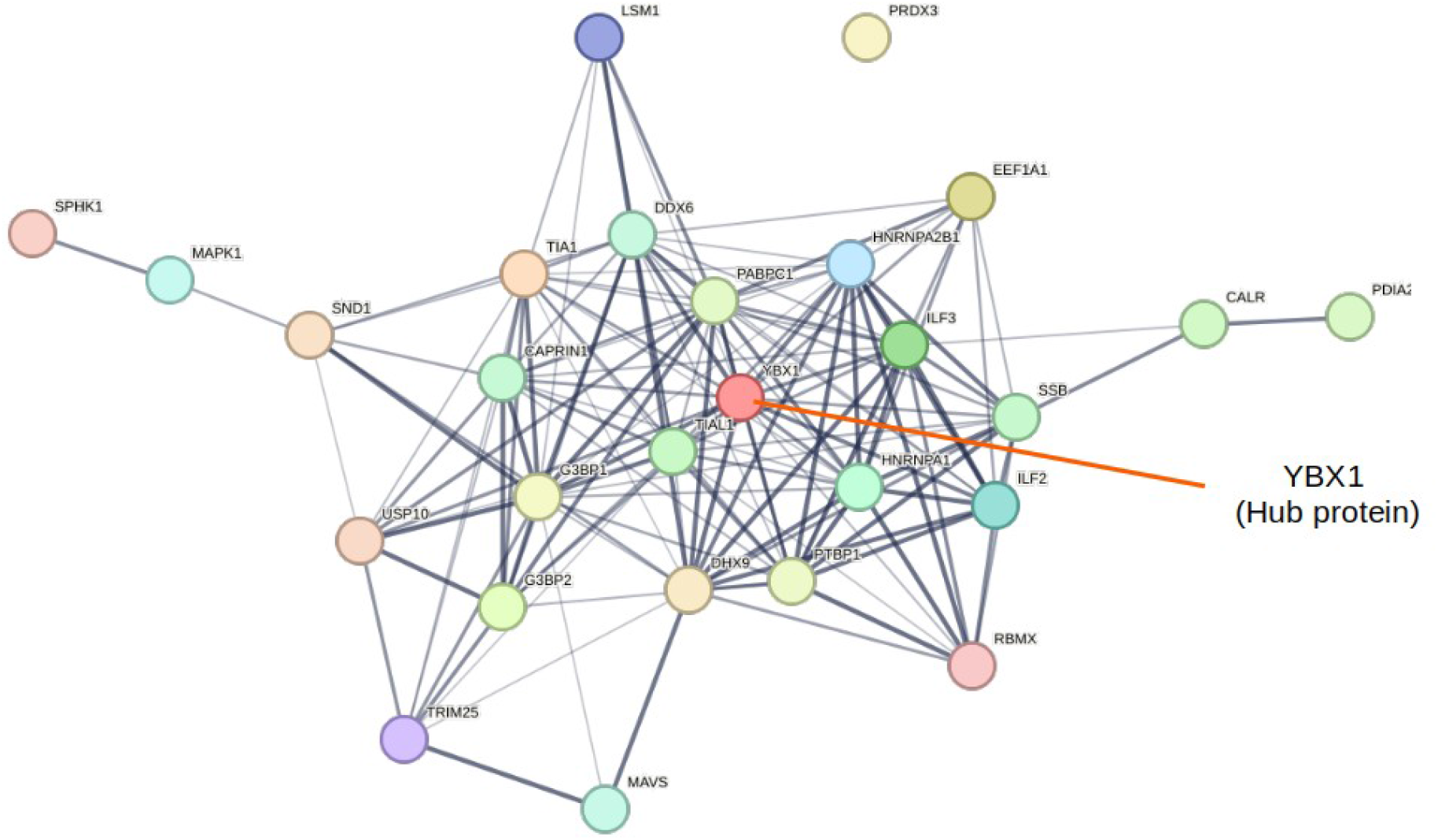
Network analysis of proteins targeted by 3’UTR domain

## Conclusion

Cross species target specificity of miRNA has been a much disputed topic yet with many evidences. In our study we tried to predict the human miRNAs that have specificity for dengue virus genome. Four of the miRNAs predicted to have cross specie specificity are not reported to target dengue genome till date. Since gene silencing small RNA is quite prevalent in nature the miRNAs can be thought of as a potential therapeutic candidates for dengue affected individuals. Inspite of the fact that hsa-miR-4758-3p_MIMAT0019904, hsa-miR-6760-3p_MIMAT0027421, hsa-miR-6839-3p_MIMAT0027581, hsa-miR-6842-3p_MIMAT0027587 can be thought of as novel therapeutic candidates we are completely aware of certain limitations of in silico study. There are evidences that cross specie targetting of miRNA might also enhance pathogenicity of an organism. Thus in vitro and in vivo analysis of the presented data should provide us with a complete insight of the predicted therapeutic strategy.

